# Neonatal surgical injury alters social behaviour and cortical network responses to adult incision

**DOI:** 10.64898/2026.09.04.749387

**Authors:** Sara Hestehave, Pi-shan Chang, Simon Beggs

## Abstract

Early-life painful events can influence brain development and behaviour later in life, yet the neural mechanisms underlying these long-term effects remain poorly understood. In particular, exposure to pain or tissue injury during early postnatal life has been associated with altered sensory processing, emotional regulation and behavioural responses later in life. Here we investigated whether early-life surgical pain alters adult behavioural and cortical network responses to injury. Using a rat model of neonatal hindpaw incision followed by repeat adult surgical incision, we combined behavioural analysis with continuous electrophysiological recordings from the medial prefrontal cortex (mPFC) and primary somatosensory cortex (S1) in freely moving animals.

Early-life incision did not produce an overt alteration in adult social behaviour under baseline conditions. However, following adult injury, rats with prior neonatal incision showed altered social behavioural responses compared with animals receiving an incision only in adulthood. At the network level, adult incision increased beta–gamma phase–amplitude coupling within the mPFC and S1–mPFC theta coherence in animals experiencing their first injury, but not in those incised neonatally. Our evidence indicates that the relationship between mPFC coupling and social interaction differed according to neonatal injury history, revealing a lasting reorganisation of cortical network recruitment and brain–behaviour coupling after repeat injury.

## Introduction

Social behaviour is dynamically adjusted according to an individual’s physiological and environmental state. In rodent models, pain, inflammation and tissue injury can transiently reduce social engagement (Bluthé et al., 2000; Baldwin et al., 2022) and this can be induced experimentally in humans (Eisenberger et al., 2010). This state-dependent change is commonly interpreted as a component of a defensive response to modify engagement with the social environment when under physiological threat. Whether this capacity to recruit these social responses is shaped by painful experiences during early life is unknown.

Early-life events play a critical role in shaping brain development and can exert long-lasting effects on social and emotional behaviour. This is clinically relevant to preterm infants who show an increased prevalence of socially withdrawn behaviour, peer-relationship difficulties and reduced participation in social relationships during adolescence and adulthood (Eryigit-Madzwamuse et al., 2015; Bilgin, 2025; Gonen et al., 2025). These outcomes reflect the combined effect of preterm birth and neonatal intensive care and the contribution made by painful and tissue-damaging treatment is unclear. Infants born preterm undergo repeated skin-breaking procedures (Laudiano-Dray et al., 2020) and have a high prevalence of surgery during neonatal intensive care (Morriss et al., 2014). These exposures occur during a period of rapid, activity-dependent maturation of sensory, cognitive and affective neural circuits. Within preterm cohorts, greater cumulative exposure to neonatal procedural pain has been associated with altered white matter, cortical and thalamocortical development (Brummelte et al., 2012; Ranger et al., 2013; Duerden et al., 2018), poorer cognitive and motor outcomes (Grunau et al., 2009; Vinall et al., 2014; Giordano et al., 2023), and increased internalising and withdrawn behaviour across childhood (Ranger et al., 2014; McLean et al., 2022). Major or repeated surgery before term equivalent age is similarly associated with poorer later developmental outcomes (Morriss et al., 2014; Gano et al., 2015). While these reports suggest that painful early-life events may influence neural circuits underlying social and emotional behaviour, clinical studies cannot separate the relative contributions of tissue injury from prematurity, illness, inflammation, anaesthesia, analgesia and the broader intensive care environment.

Pre-clinical models provide an important means of determining how early surgical injury itself influences developing neural circuits and subsequent behaviour. Rodent models of neonatal incision show persistent alterations in nociceptive processing and in the organisation of developing sensory networks (Ren et al., 2004; Walker et al., 2009; Beggs et al., 2012; Moriarty et al., 2019; Chang et al., 2022). To date, the majority of research has concentrated on the sensory consequences of neonatal injury, and it remains unclear whether these effects extend to those circuits governing social behaviour. It therefore remains unclear whether neonatal surgical injury alters social behaviour directly or instead changes how social engagement is regulated during a later injury.

The medial prefrontal cortex (mPFC) integrates sensory, cognitive and emotional information to guide adaptive behavioural responses(Ong et al., 2019). The primary somatosensory cortex (S1) contributes to the representation of nociceptive sensory information, and functional interactions between S1 and mPFC may integrate sensory input with the affective, motivational and social behavioural responses to injury (Chang et al., 2016, 2020, 2022; Tan and Kuner, 2021). Communication within these regions is supported by coordinated oscillatory activity. Phase-amplitude coupling coordinates neural activity across temporal scales within a region, whereas inter-regional coherence provides a measure of functional coupling between cortical networks. This circuit activity undergoes prolonged postnatal development and therefore particularly sensitive to early life perturbations (Chang et al., 2016, 2020, 2022). Whether these effects persist into adulthood and how they influence network level responses to subsequent injury are not known. In particular, it is unknown whether early-life surgical injury alters the ability of prefrontal circuits to engage protective or adaptive network states following further painful events such as surgical injury in adulthood.

Here, we use combined behavioural and electrophysiological analysis of functional connectivity and cross-frequency coupling to test whether neonatal surgical injury produces a latent alteration in the regulation of adult social behaviour and identify the cortical network changes associated with it. Using a rat model of neonatal hindpaw incision followed by incision in adulthood, we assessed social behaviour tests with continuous wireless telemetric recordings of neural activity from the mPFC and S1 cortex in untethered, freely moving animals. We tested whether neonatal injury altered social approach and social novelty and whether these behavioural responses were associated with changes in mPFC beta-gamma phase-amplitude coupling and S1-mPFC functional connectivity. Neonatal incision did not produce an overt alteration in adult social behaviour under baseline conditions but modified social approach following adult incision. At the network level, prior neonatal incision was associated with the absence of injury-linked increases in mPFC coupling and S1-mPFC coherence observed in animals receiving adult incision only, consistent with impaired engagement of adaptive network states. Together, our findings reveal a mechanism through which early-life sensory experiences reprogramme adult prefrontal circuit activity, revealing a latent vulnerability that only emerges under challenge.

## Materials and methods

### Experimental animals

All experiments were performed in accordance with the United Kingdom Animal (Scientific Procedures) Act 1986. Reporting is based on the ARRIVE Guidelines for Reporting Animal Research developed by the National Centre for Replacement, Refinement and Reduction of Animals in Research, London, United Kingdom. Male Sprague-Dawley rat pups were obtained from the Biological Services Unit, University College London. Rats were housed in cages of five age-matched animals (>P21) or with the dam and littermates (P) 3 to 21 under controlled environmental conditions (24–25°C; 50–60% humidity; 12 h light/dark cycle) with free access to food and water. In the case of rat pups, handling and maternal separation were kept to a minimum. All animals were exposed to the same standard caging, handling and diet throughout development.

### Plantar hind-paw incision

Male rat pups were anaesthetized (2% isoflurane in 100% oxygen) and plantar hindpaw incision performed on postnatal Day 3 (P3) as described previously(Walker et al., 2009; Chang et al., 2016). Briefly, a midline longitudinal incision was made through the skin and fascia extending from the midpoint of the heel to the proximal border of the first footpad and the underlying plantar muscle elevated and incised. The same relative length of incision was performed in adult animals. Skin edges were closed with 5–0 nylon suture (Ethicon). After plantar hindpaw incision, rats were placed in a recovery chamber and allowed to recover from the general anaesthesia before returning to their home cage.

Four experimental groups were used: (**I** = hindpaw incision; **N** = no incision but equivalent anaesthesia, handling and maternal separation):

**II:** Neonatal incision and repeat incision in adulthood.

**NI:** Adult incision only

**IN:** Neonatal incision only

**NN:** Control

### Surgical Preparation and Transmitter Implantation for Long-term Recording

Rats were anaesthetised with 2.5-3 % isoflurane in 100% oxygen (flow rate of 1-1.5 litre/min). Body temperature was maintained with a heat blanket during surgery. A transmitter (A3028D-DDA, Open Source Instruments, Brandeis, Boston, USA) was implanted subcutaneously with the depth recording electrodes (J-electrode (wire 125-μm diameter 316SS 10kOhm impedance), a Teflon-insulated stainless steel electrode, Open Source Instruments, Brandeis, Boston, USA) positioned in mPFC (3.2 mm anterior, 0.5 mm lateral, 4 mm ventral) and primary somatosensory hindpaw cortex (1 mm posterior, 2.5 mm lateral, 2 mm ventral) (Chang et al., 2016). The reference electrode was implanted over the cerebellum posterior to lambda. The whole assembly was held in place with dental cement (Simplex Rapid, Acrylic Denture Polymer, UK). A subcutaneous injection of bupivacaine and metacam was provided for post-surgical pain management. At the end of surgery, enrofloxacin (5mg/kg, Baytril, Bayer health care) was administered subcutaneously. The animals were placed in a temperature controlled (25°C) recovery chamber until ambulatory and closely monitored at least 1-2 hours before returning to their home cage to allow recovery for at least 14 days after surgery.

Continuous LFP recordings were made during recording sessions (bandpass filter: 0.2 Hz to 160 Hz, 512Hz sampling rate with 16 bit resolution) using LWDAQ Software (Open Source Instruments, Brandeis, Boston, USA). At the end of the experiments, animals were euthanized and brains collected to check the histological location of the electrode track. This procedure allowed us to verify recording electrode locations, and LFP data were only included in the study if electrode tips were located in mPFC and S1

### Social behaviour

#### Three-chambered social approach tests

The three-chambered test was used to assess social approach and social novelty preference in adult rats (Silverman et al., 2010). The apparatus consists of an open-topped acrylic box (120cm L x80cm W x40cm H) divided into three chambers with two opaque acrylic walls. Dividing walls had retractable doorways allowing access into each chamber. Test rats were confined in the centre chamber at the beginning of each trial. At the start of each 10-min phase, the test rat was placed in the centre chamber and then allowed free access to all chambers. Two identical wire cages were positioned in the side chambers. During the habituation phase (phase 1), each of the two side chambers contained an empty cage. During the sociability phase (phase 2), a stranger rat was enclosed in one of the cages in a side chamber. During the social novelty phase, a new stranger rat and now familiar initial stranger were enclosed in the cages. Exploration of an enclosed rat or a cage is defined as when a test rat is oriented toward the grid cage with the distance between the nose and the cage less than 1 cm. The time spent exploring enclosed rats or empty cages was recorded by a camera mounted overhead and analyzed by an automated tracking program (AnyMaze). All the stranger rats were habituated to being enclosed in a cage in the three-chamber apparatus prior to the experiment.

Time spent investigating each cage was quantified using automated tracking software (Ethovision XT, Noldus).

S**ociability index** was calculated as:

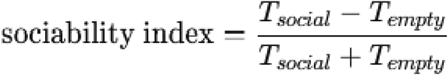

where *T*_*social*_ represents time spent investigating the cage containing the stimulus rat.

### Analysis of electrophysiology data

Data analysis was performed with Brainstorm (http://neuroimage.usc.edu/brainstorm) and custom Matlab scripts (The Mathworks Inc.MA, USA).

### Resting-LFP data processing

#### Pre-processing

Continuous LFP recordings from each region were segmented into 100s epochs. Each epoch was visually inspected for artefacts prior to further analysis. Any epochs that exhibited artefacts during visual inspection were excluded from subsequent analysis.

#### Phase-amplitude Coupling Analysis

PAC is calculated between the phase of a low frequency activity and the amplitude of a high frequency activity to determine if the phase of the lower frequency modulates the amplitude of the higher. The signal in each epoch (100s) was filtered into a range of frequency pairs, with centre frequencies for the low frequencies ranging from 2-30 Hz, and for the high frequencies ranging from 60–90 Hz. Subsequently, Phase-amplitude coupling was assessed between each low-frequency and high-frequency filtered signal.

#### Coherence analysis

Coherence analysis of the LFP data was performed to detect cortical connectivity patterns pre and post-incision. The complex coherence between S1 and mPFC signals was calculated as the cross-spectrum between the signals and normalized by the square root of the power spectrum product of the two signals. Given that coherence is a normalized measure of the correlation between two signals its amplitude can vary from 0 to 1, where 0 means that the frequency components of both signals are linearly independent, while 1 means the frequency components of the two signals give the maximum linear correlation.

### Quantification and Statistical Analysis

Statistical analyses were performed using GraphPad Prism 10 (GraphPad Software) and IBM SPSS Statistics (IBM). Data are presented as mean ± SEM or as mean differences with 95% confidence intervals (CIs), as indicated. Normality was assessed using the Shapiro–Wilk test where applicable. Statistical significance was defined as P < 0.05.

For the three-chamber test, time spent investigating the two cages within each phase was compared within each experimental group using two-tailed paired Student’s t-tests. These within-group comparisons are reported as unadjusted P values. To test whether behavioural preference differed according to injury history, an animal-level preference score was calculated as the paired difference in investigation time between the two cages. Preference scores were analysed using a two-way ANOVA of neonatal incision, adult incision and their interaction as between-subject factors.

For electrophysiological analyses of responses to adult incision, NN and IN animals were combined into a single control group. This pooling was not pre-specified but followed an interim analysis in which no detectable differences between NN and IN animals were observed for the relevant electrophysiological outcomes. Phase-amplitude and theta coherence proportional changes following manipulation (D0/Pre ratios) were compared using ordinary one-way ANOVA. Recordings were obtained in home cages under identical environmental conditions, and long epoch averaging with artefact rejection reduced state-related variance. PAC and coherence are state dependent, and epochs were not annotated for locomotor state; thus residual differences in D0 state composition may influence coupling measures. Normalised phase–amplitude coupling values were compared using ordinary one-way ANOVA followed by Bonferroni-adjusted pairwise comparisons.

Between-group differences in theta coherence were additionally quantified using estimation statistics. Unpaired mean differences between groups were calculated with bias-corrected and 95% CIs obtained from 5,000 bootstrap resamples. Two-sided permutation P values were calculated.

Associations between mPFC beta–gamma phase–amplitude coupling and the social-interaction index were examined in NI and II animals using ordinary least-squares linear regression. Regression slopes were compared using analysis of covariance, incorporating group, social-interaction index and their interaction in the model.

## Results

### Early-life surgical pain does not produce overt changes in adult social behaviour

We first asked whether neonatal surgical incision altered adult social behaviour in the absence of further injury. During habituation, neither naïve control rats (NN; P = 0.822) or rats exposed to neonatal incision alone (IN; P = 0.158) showed a significant preference for either chamber (Figure 2B). Baseline behavioural profiling showed no significant differences in mean speed, maximum speed, time immobile or immobile episodes between groups (data not shown).

**Figure 1.**
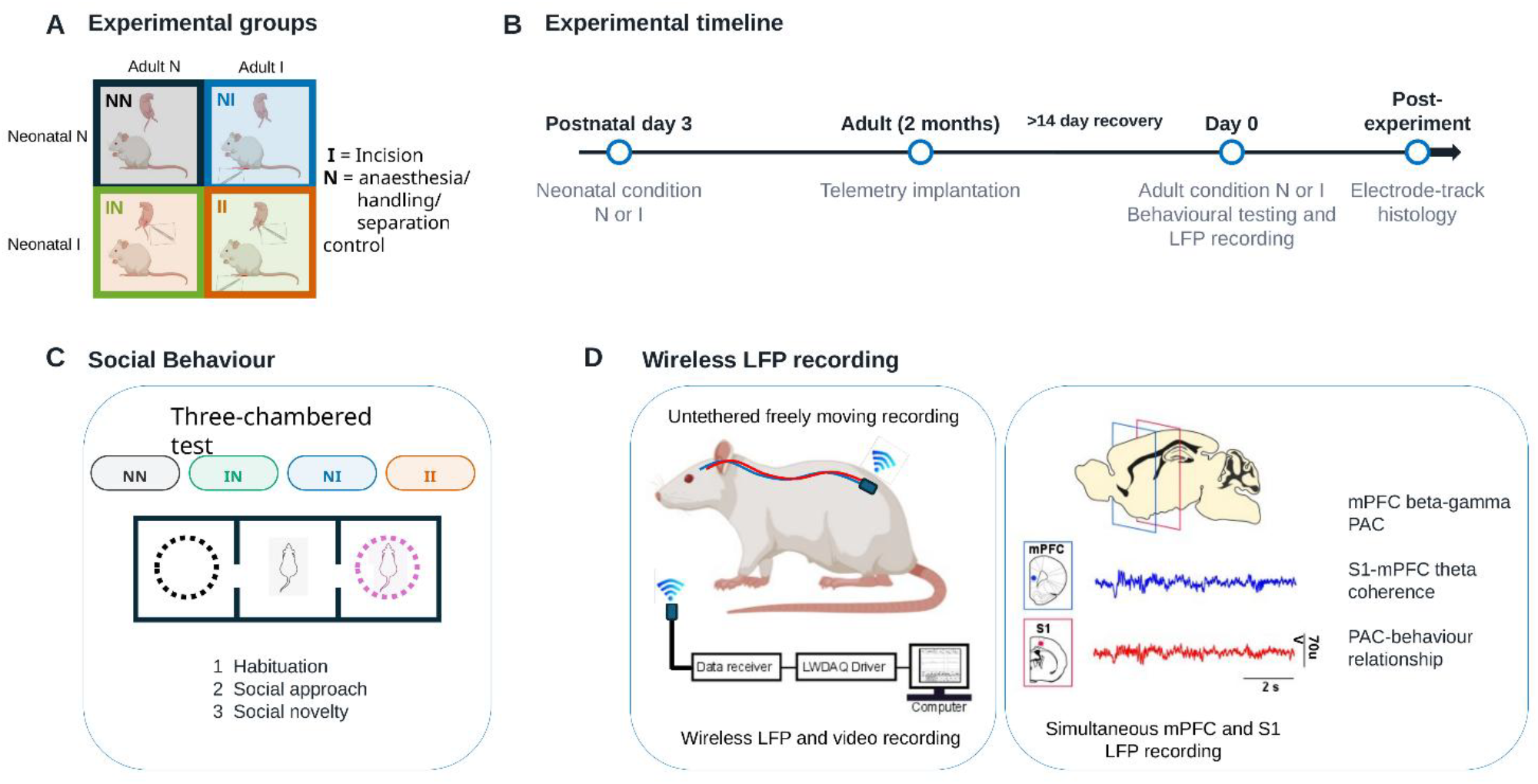
Experimental design for simultaneous behavioural and cortical recording. (A) Four experimental groups were defined by neonatal and adult treatment. The first letter denotes the neonatal condition and the second the adult condition: NN, no incision at either age; IN, neonatal incision only; NI, adult incision only; and II, neonatal and adult incision. N denotes no incision with equivalent anaesthesia, handling and separation controls; I denotes plantar hindpaw incision. (B) Rats underwent neonatal incision at P3, wireless transmitter and electrode implantation at 2 months of age, and adult incision followed by behavioural testing and recording on Day 0 (D0). Electrode locations were verified histologically after the experiment. (C) Three-chamber testing comprised habituation, social approach and social novelty. (D) Untethered local field potentials were recorded simultaneously from the medial prefrontal cortex (mPFC) and primary somatosensory cortex (S1). Analyses quantified mPFC beta–gamma phase–amplitude coupling (PAC), S1–mPFC theta-band coherence and the relationship between PAC and social behaviour.

**Figure 2.**
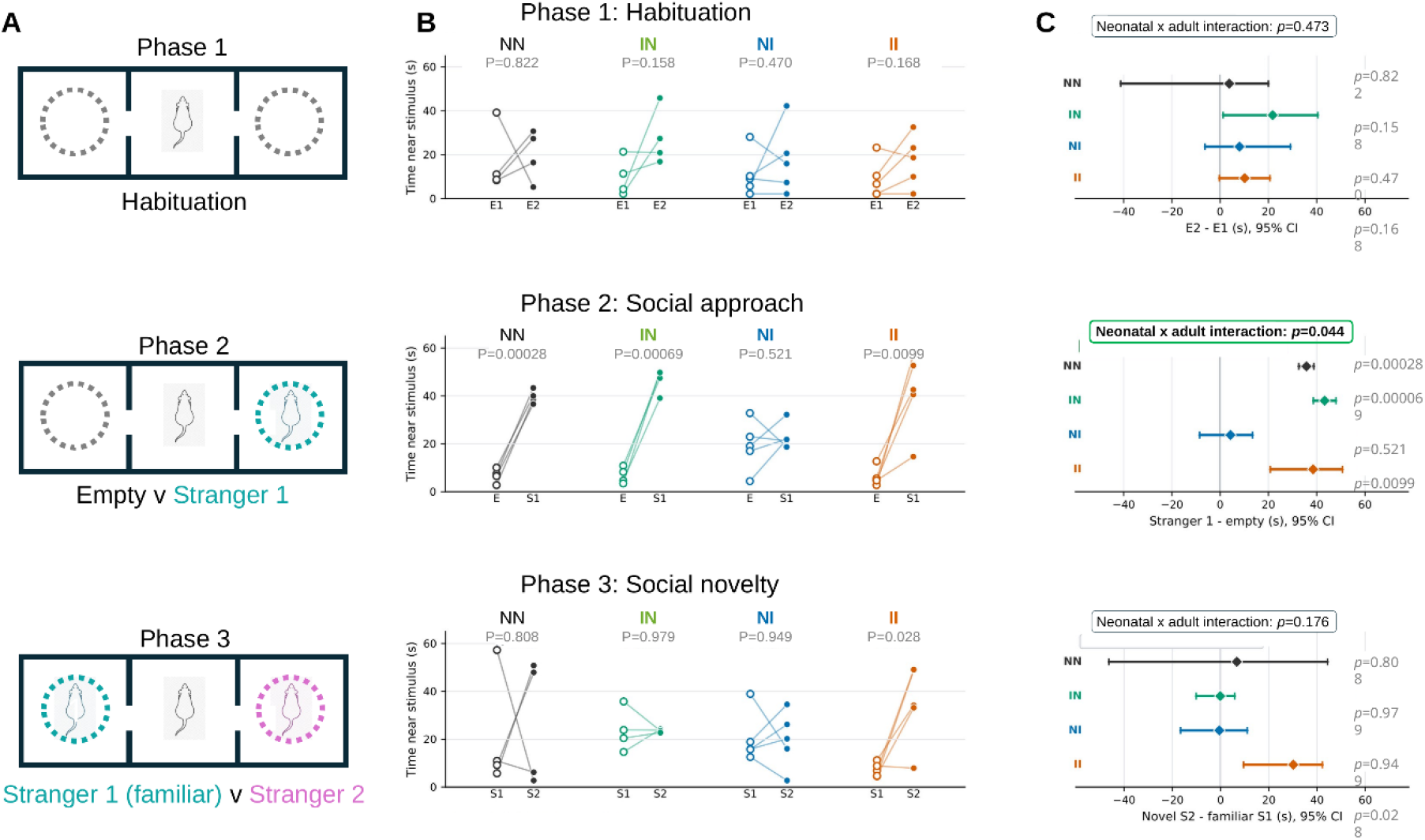
Early-life surgical pain alters social behaviour following adult injury. (A) Three-chamber test sequence. During habituation, both stimulus cages were empty (E1 and E2). During social approach, animals chose between an empty cage (E) and an unfamiliar conspecific (Stranger 1; S1). During social novelty, animals chose between the now-familiar S1 and a newly introduced unfamiliar conspecific (Stranger 2; S2). (B) Time spent near each stimulus. Open and filled symbols denote the first and second stimulus labels, respectively; lines connect observations from the same animal. (C) Mean paired difference in investigation time with 95% confidence interval (CI). NN and IN, n = 4 per group; NI and II, n = 5 per group. Within-group t-tests across phases were descriptive and unadjusted; effect sizes are shown with confidence intervals. Interaction P values were 0.473 for habituation, 0.0444 for social approach and 0.176 for social novelty. Exact within-group P values are shown in the panels; P < 0.05 was considered significant.

During the social-approach phase (Phase 2), both NN and IN animals spent significantly more time near Stranger 1 than the empty cage (NN, P = 0.00028; IN, P = 0.00069). During the subsequent social-novelty phase (Phase 3), neither group showed a significant preference between the novel Stranger 2 and the now-familiar Stranger 1 (NN, P = 0.808; IN, P = 0.979). Thus, NN and IN animals displayed the same overall behavioural profile of intact social approach but no significant preference for social novelty. Neonatal incision alone therefore did not produce an overt alteration in adult social behaviour.

### Neonatal incision modifies social responses following adult injury

We next examined whether neonatal injury history influenced social behaviour following adult incision. During habituation, neither animals receiving their first incision in adulthood (NI; P = 0.470) nor animals exposed to both neonatal and adult incision (II; P = 0.168) showed a cage-location preference. There was no neonatal incision × adult incision interaction during this phase (P = 0.473; Figure 2C).

During the social-approach phase, NI animals did not preferentially spend time near Stranger 1 rather than the empty cage (P = 0.521). In contrast, II animals retained a significant preference for Stranger 1 (P = 0.0099; Figure 2B). Factorial analysis of the social-approach difference scores demonstrated a significant neonatal incision × adult incision interaction (P = 0.0444; Figure 2C). This interaction suggests a dependence of social approach on neonatal injury history, with social approach absent in rats experiencing their first incision in adulthood but retained following repeat incision in animals previously exposed to neonatal injury.

In the subsequent social novelty phase, NI animals showed no preference between the novel Stranger 2 and the familiar Stranger 1 (P = 0.949), whereas II animals spent significantly more time near Stranger 2 (P = 0.0281; Figure 2B). This result therefore shows a significant social-novelty preference within the II group. However, the neonatal incision × adult incision interaction did not reach statistical significance for social novelty (P = 0.176; Figure 2C) and therefore does not establish a statistically significant difference in novelty preference between injury groups.

Together, these findings indicate that neonatal incision alone does not cause an observable social deficit in adulthood. Instead, neonatal injury modifies the behavioural response recruited following subsequent adult injury, with a significant effect on social approach and an additional within-group social-novelty preference following repeat incision.

Having established that neonatal incision modified social behaviour following adult injury, we next asked whether this effect was accompanied by altered recruitment of cortical networks linking nociceptive processing with behaviour. No significant differences were detected between NN and IN animals and were therefore combined into a single Control group for subsequent electrophysiological analyses.

### Early-life surgical pain alters prefrontal phase–amplitude coupling during adult injury

Group-averaged comodulograms indicated differences in beta–gamma phase– amplitude coupling (PAC) within the mPFC following adult incision (Figure 3A,B). Analysis of the modulation index confirmed a significant group effect in the mPFC (one-way ANOVA, F(2,28) = 5.663, P = 0.0086; Figure 3D). PAC was higher in NI than Control animals (mean difference = 0.277, 95% CI 0.047–0.506; Bonferroni-adjusted P = 0.0140) and II animals (II − NI = −0.296, 95% CI −0.558 to −0.034; P = 0.0226). In contrast, II animals did not differ from Control (mean difference = −0.020, 95% CI −0.249 to 0.210; P > 0.9999). Thus, the mPFC beta–gamma coupling increase on D0 seen in NI animals was absent in animals with a prior neonatal incision.

**Figure 3.**
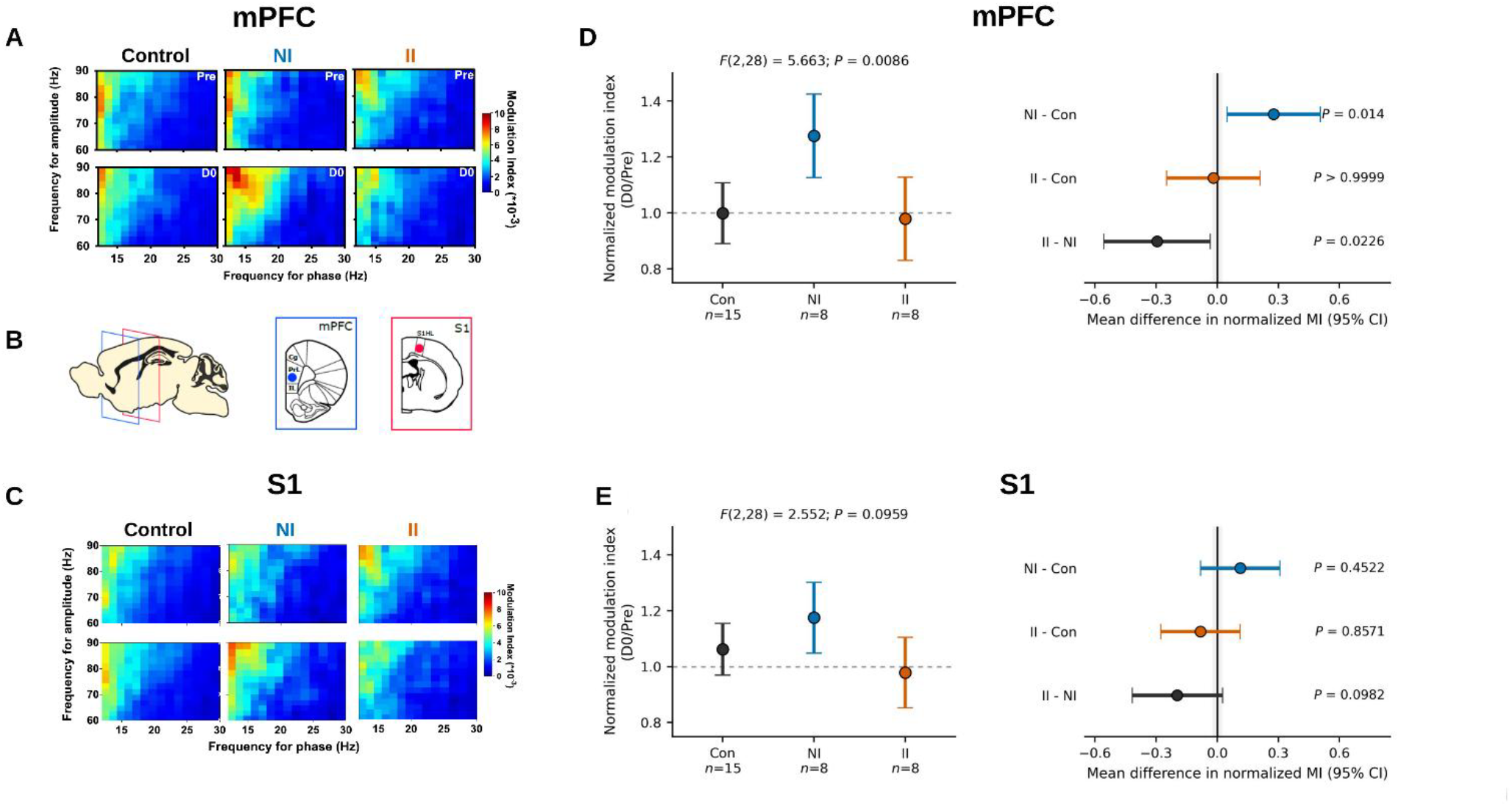
Neonatal incision alters phase–amplitude coupling in medial prefrontal cortex following adult injury. (A) Group-averaged mPFC phase–amplitude comodulograms before the adult condition (Pre) and on Day 0 (D0) for Control, NI and II rats. (B) Recording locations in mPFC and S1. (C) Corresponding S1 comodulograms. (D) Left, mPFC modulation index at D0 normalised to Pre; points and error bars show group means and 95% CIs. Right, pairwise mean differences with 95% CIs. (E) Corresponding S1 analysis. Control, n = 15; NI, n = 8; II, n = 8. Control comprises NN and IN animals. Normalised modulation indices were compared by one-way ANOVA followed by Bonferroni-adjusted multiple comparisons.

This effect was not observed in S1 (Figure 3C, E). Normalised S1 beta–gamma coupling did not differ significantly between Control, NI and II animals (one-way ANOVA, F(2,28) = 2.552, P = 0.0959; Figure 3E), and all Bonferroni-adjusted pairwise comparisons were non-significant (P ≥ 0.0982). The effect of neonatal injury history on adult injury-associated PAC was therefore specific to the mPFC.

### Early-life surgical pain alters prefrontal–somatosensory functional connectivity following adult injury

We next investigated whether neonatal incision also modified long-range cortical communication by assessing functional coupling between S1 and mPFC. We examined coherence across the full frequency spectrum before (Pre) and immediately after adult incision (D0; Figure 4A). We then quantified mean coherence across the theta band (4-8 Hz), indicated by the grey shading and expressed this is as proportional change (D0/Pre; Figure 4B). Theta coherence in NI animals had a greater proportional increase (D0/Pre) than in control animals (mean difference = 0.135, 95% CI 0.060–0.227; permutation P = 0.0088; Figure 4B). II animals did not differ from Control animals (mean difference = 0.026, 95% CI −0.047 to 0.098; P = 0.5470) but had lower proportional change in coherence than NI animals (II − NI = −0.108, 95% CI −0.201 to −0.035; P = 0.0284). Thus, as observed for mPFC PAC, the larger proportional change in S1–mPFC theta coherence associated with adult incision was absent in animals previously exposed to neonatal incision. These findings indicate that neonatal injury history modifies both local prefrontal network dynamics and long-range cortical coordination during subsequent injury.

**Figure 4.**
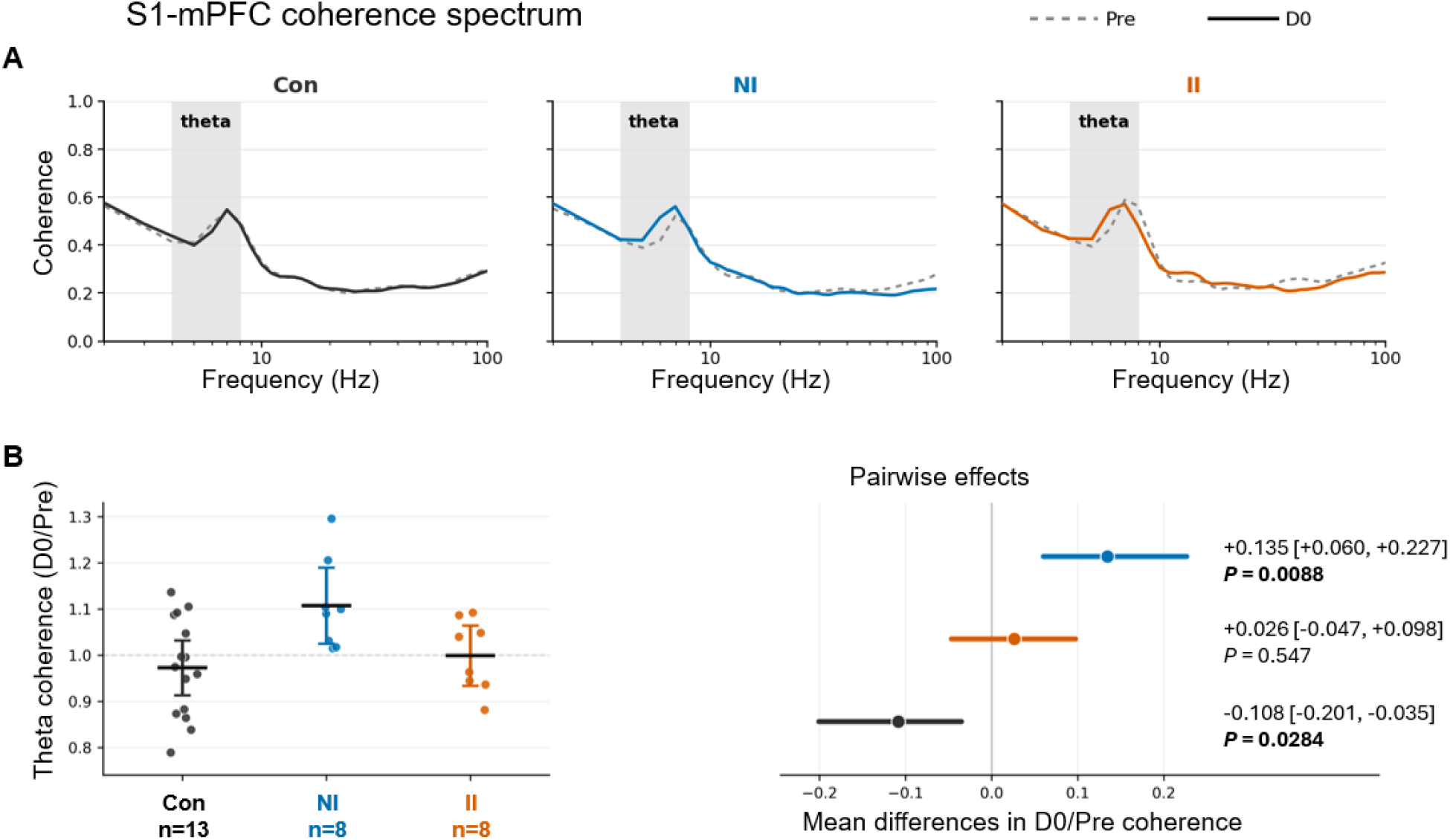
Neonatal incision attenuates adult injury-evoked changes in S1-mPFC theta coherence. (A) S1–mPFC coherence spectra across the theta band before (Pre; grey dashed lines) and immediately after adult manipulation (D0; solid lines) in control, NI and II rats. Grey shading indicates the theta frequency range (4-8 Hz) used for quantification. (B) Left, individual-animal theta coherence proportional change; horizontal bars show group means and 95% CIs. Right, pairwise mean differences in normalised coherence with bootstrap 95% CIs. Control, n = 15; NI, n = 8; II, n = 8. Control comprises NN and IN animals.

### Neonatal injury alters the relationship between prefrontal cortex coupling and social behaviour in adulthood

We next examined whether individual differences in mPFC beta–gamma PAC were associated with social behaviour following adult incision. The coupling-behaviour analysis was time-locked to the social testing periods. The relationship between PAC and social-interaction index differed significantly between NI and II animals (comparison of regression slopes: F(1,7) = 19.18, P = 0.0032; Figure 5). The relatively small numbers of animals contributing to the regressions also means that the precise slope estimates should be interpreted cautiously.

**Figure 5.**
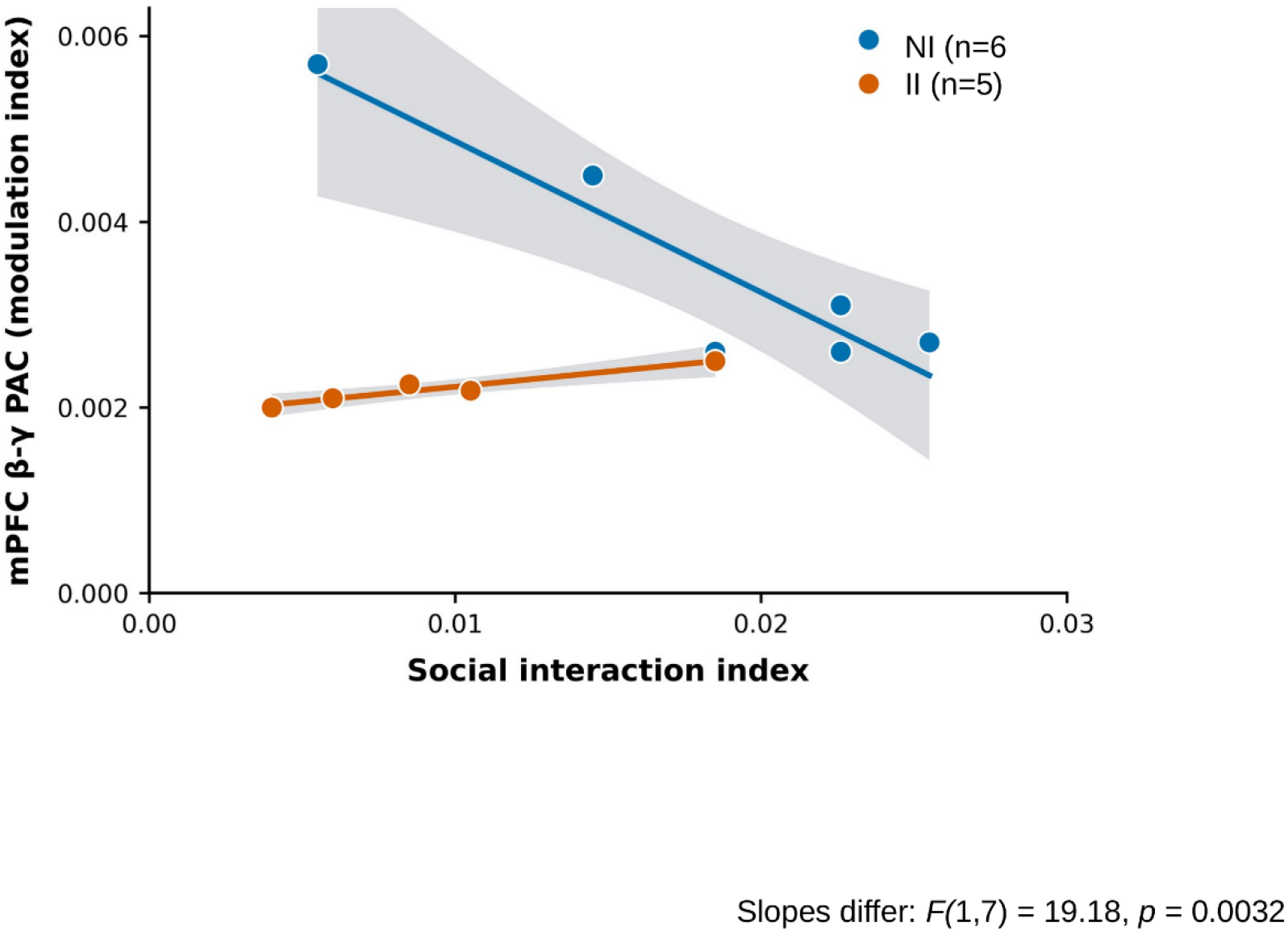
Neonatal incision alters the relationship between medial prefrontal beta– gamma phase–amplitude coupling and social interaction following adult injury. Relationship between the social interaction index (interaction time/total time) and the mPFC beta– gamma PAC modulation index at D0 in animals receiving adult incision. Each point represents one animal; lines show ordinary least-squares linear regressions and grey shaded regions show 95% CIs. NI rats showed a negative association (slope = −0.1628, 95% CI −0.2550 to −0.0706; R^2^ = 0.857, P = 0.0080; n = 6), whereas II rats showed a positive association (slope = +0.0325, 95% CI +0.0160 to +0.0490; R^2^ = 0.929, P = 0.0081; n = 5). ANCOVA testing the group × social interaction term (equivalent to an extra-sum-of-squares F test) showed that the slopes differed between groups, F(1, 7) = 19.18, P = 0.0032.

In NI animals, mPFC PAC was negatively associated with social interaction (slope = −0.1628, 95% CI −0.2550 to −0.0706; R^2^ = 0.857, P = 0.0080). In contrast, II animals showed a positive association between PAC and social interaction (slope = 0.0325, 95% CI 0.0160–0.0490; R^2^ = 0.929, P = 0.0081). The difference between the slopes was 0.1953 (II − NI; 95% CI 0.0898–0.3007), demonstrating that neonatal injury altered the direction of the relationship between prefrontal network dynamics and social behaviour following adult incision.

Together, these findings show that neonatal incision not only prevented the increases in mPFC beta–gamma PAC and S1–mPFC theta coherence changes normally recruited during surgical incision injury in adulthood, but also changed how prefrontal coupling related to behavioural responses across individual animals.

## Discussion

Early-life experiences can exert lasting influences on neural circuit development and behaviour. Here, neonatal surgical injury did not alter adult social behaviour under baseline conditions but modified the social response to a later incision. This behavioural effect was accompanied by altered recruitment of local and long-range cortical networks and by a reversal in the relationship between mPFC coupling and social interaction.

Transient reductions in social interaction occur following injury, infection or inflammation and are commonly interpreted as components of defensive or sickness-related behavioural states (Bluthé et al., 2000; Eisenberger et al., 2010; Baldwin et al., 2022). Such changes may reduce engagement with potentially threatening stimuli and redirect behaviour towards recovery. The absence of social-approach preference in NI animals may therefore form part of the adaptive behavioural response to acute injury rather than a general impairment of sociability. In contrast, animals exposed to both neonatal and adult incision retained a significant preference for social approach, indicating that neonatal injury history modified the social response to adult incision. II animals also showed a within-group preference for the novel stranger, whereas NI animals did not. However, because the neonatal incision x adult incision interaction was not significant for social novelty, this observation should be treated with caution.

Pain is a multidimensional experience involving sensory, affective and motivational processes, and injury can alter motivation and willingness to engage with unfamiliar conspecifics (Low and Fitzgerald, 2012; Baldwin et al., 2022). The present behavioural effects may therefore reflect changes in the affective or motivational response to injury rather than differences in sensory processing alone. Through its integration of nociceptive, affective and social information, the mPFC is well placed to contribute to this context-dependent regulation of behaviour (Franklin et al., 2017; Murugan et al., 2017; Ong et al., 2019). Adult incision was associated with increased mPFC beta-gamma PAC in NI animals, whereas PAC in II animals remained comparable with controls. Because corresponding group differences were not detected in S1, this effect was regionally selective. The increase in NI animals may reflect recruitment of local prefrontal integration during the evaluation of injury and organisation of an appropriate behavioural response. Its absence following neonatal incision suggests that early injury alters how this prefrontal network state is recruited during later challenge. Neonatal incicion similarly modified long-range cortical coordination. Animals experiencing their first incision in adulthood showed S1 and mPFC theta coherence, whereas animals previously exposed to neonatal incision remained comparable with controls. Theta sunchronisation is widely implicated in long-range coordination between distributed neural populations (Jones and Wilson, 2005). In the present context, increased S1-mPFC theta coherence may therefore reflect coordination of sensory and prefrontal processing during acute injury. Its absence in II animals indicates altered recruitment of this injury-associated network response rather than an absolute loss of baseline functional connectivity.

The relationship between mPFC PAC and social interaction also depended on neonatal injury history. In NI animals, greater mPFC beta-gamma PAC was associated with lower social interaction, whereas the relationship was positive in II animals. These differing associateions suggest that neonatal injury changes how prefrontal network states map onto behavioural output during adult injury, rather than merely shifting the overall magnitude of cortical coupling. However, the small numbers contributing to these regressions requires cautious interpretation and replication in a larger cohort. These context-dependent effects are consistent with developmental programming. The mPFC undergoes marked postnatal maturation in neuronal properties and in the balance of excitatory and inhibitory synaptic inout, with synaptic refinement continuing through adolescence (Drzewiecki et al., 2016; Kroon et al., 2019). Early nociceptive input could therefore modify the developing organisation of prefrontal networks and their subsequent recruitment during injury.

These findings extend previous evidence for persistent changes in spinal nociceptive processing and developing sensory pathways (Ren et al., 2004; Walker et al., 2009, 2015; Beggs et al., 2012; Moriarty et al., 2019; Chang et al., 2022), by showing altered recruitment of cortical networks that integrate nociceptive information with behaviour. They identify prefrontal network flexibility as an additional potential locus of developmental programming by early-life injury. This principle has translational relevance to early-life medical care. Infants born prematurely or undergoing surgery experience repeated nociceptive events during critical periods of brain maturation. These findings are relevant to infants born very preterm or requiring neonatal surgery, who undergo repeated nociceptive events during periods of rapid brain development. Neonatal pain-related stress and surgery are associated with later differences in somatosensory, emotional and cognitive outcomes (Gano et al., 2015; Walker et al., 2018a, 2018b; McLean et al., 2022; Giordano et al., 2023). Although direct translation from this rodent model requires caution, the present findings suggest a circuit-level framework through which early nociceptive experience could modify responses to later injury or stress. Such effects may remain clinically dormant until the relevant systems are challenged, conceptually consistent with latent vulnerability (McCrory and Viding, 2015).

Several limitations should be acknowledged. Firstly, the study was conducted in male rats, and it remains to be determined whether equivalent programming effects occur in females. Secondly, the regression analyses involved relatively small groups and their correlational nature requires caution in any conclusions about causality. Finally, LFP recordings describe population-level network activity but do not identify the cellular, synaptic or molecular mechanisms through which neonatal incision modifies circuit development. Defining these mechanisms and determining whether the altered network responses can be restored will be important objectives for future work.

## Conclusions

Early-life surgical pain altered the behavioural and cortical network responses recruited during surgical injury in adulthood. Although neonatal incision alone did not produce an overt deficit in adult social behaviour, it modified social responses to adult incision, was associated with the absence of injury-linked increases in mPFC beta-gamma coupling and S1-mPFC theta coherence, and changed the relationship between prefrontal coupling and social interaction. These findings identify neonatal injury as a developmental programming event that reorganises brain-behaviour coupling and influences how cortical networks respond to a later challenge.

